# A Simple Cross-Linking/Mass Spectrometry Workflow to Study System-Wide Protein Interactions

**DOI:** 10.1101/524314

**Authors:** Michael Götze, Claudio Iacobucci, Christian Ihling, Andrea Sinz

## Abstract

We present a cross-linking/mass spectrometry (XLMS) workflow for performing proteome-wide cross-linking analyses within one week. The workflow is based on the commercially available MS-cleavable cross-linker disuccinimidyl dibutyric urea (DSBU) and can be employed by every lab having access to a mass spectrometer with tandem MS capabilities. We provide an updated version 2.0 of the freeware software tool MeroX, available at www.StavroX.com, that allows conducting fully automated and reliable studies delivering insights into protein-protein interaction networks and protein conformations at the proteome level. We exemplify our optimized workflow for mapping protein-protein interaction networks in *Drosophila* melanogaster embryos on a system-wide level. From cross-linked *Drosophila* embryo extracts, we detected 18,037 cross-link spectrum matches corresponding to 5,129 unique cross-linked residues in biological triplicate experiments at 5% FDR (3,098 at 1% FDR). Among these, 1,237 interprotein cross-linking sites were identified that contain valuable information on protein-protein interactions. The remaining 3,892 intra-protein cross-links yield information on conformational changes of proteins in their cellular environment.

## INTRODUCTION

Cells and tissues possess complex organizations that cannot fully be captured via in vitro experiments using isolated proteins. The classical techniques for high resolution protein structural analysis, such as NMR spectroscopy, X-ray crystallography, and cryo-electron microscopy (cryoEM), are usually employed for in vitro samples and might not accurately reflect the protein structures in a cell or a tissue. Studying protein-protein interactions adds an extra level of sample complexity, given that the majority of biological processes depend on hundreds to thousands of macromolecular interactions. The resulting protein networks comprise protein complexes that are organized in higher-order functional communities. Therefore, disentangling the protein interactome in cellulo presents extraordinary challenges for elucidating the functional organization of proteins.

Yeast two-hybrid (Y2H) screening^1^, co-immunoprecipitation^2^, affinity purification coupled with mass spectrometry (APMS)^3-7^, and proximity-dependent biotin identification (BioID)^8-10^ are currently being used for mapping proteinprotein interactions in the cellular environment. While these approaches are able to decipher protein networks, they do not provide information on the proteins’ morphology. Time-consuming modifications of the bait protein are required that might eventually alter the structures of native protein complexes or lose transient interaction partners. Additionally Y2H method may generate false positives due to non-specific interactions.^11^

Chemical cross-linking/mass spectrometry (XLMS) has emerged as an alternative, powerful tool for the 3D-structure analysis of proteins and protein complexes and is becoming increasingly popular in structural biology.^12-23^ The cross-linker has a defined length and imposes a distance constraint between the amino acids connected. The cross-linked proteins and protein complexes are enzymatically digested and analyzed by high-resolution mass spectrometry (MS). This permits the identification of the cross-linked amino acids yielding conclusions on their spatial proximity in the protein assembly. The resulting map of amino acid distances can be computationally reprocessed resulting 3D-structural models of the protein or protein assembly.^24-29^ The main advantage of XLMS over other protein structural techniques is that it creates comprehensive snapshots of the protein landscape with the minimum interference. XLMS experiments can be conducted within a few days making XLMS a highly attractive approach that complements existing high-resolution protein structural techniques, such as NMR spectroscopy, X-ray crystallography, and cryo-EM.^30-36^ On top of that, XLMS only requires minute protein amounts and is even applicable to intact cells^37, 38^. System-wide XLMS offers two key benefits: i) it allows capturing system-wide protein interactions for a comprehensive understanding of cellular signaling pathways and ii) it allows analyzing the conformation and the interaction of proteins in their native environment. Therefore, XLMS is currently one of the most promising MS-based approaches to derive 3Dstructural information on very large and transient protein complexes^33, 39-43^ as well as on intrinsically disordered proteins^44^.

The application of XLMS to highly complex protein mixtures, such as cell lysates^45, 46^, organelles^39, 47, 48^, intact cells^37, 38, 49^, and tissues^50^, was pioneered by Bruce^51^ in 2005. Since then, the number of research groups that have performed systemwide XLMS studies has remained limited^52^, while XLMS investigation of isolated protein and protein complexes is blossoming. Fast and reliable XLMS studies of complex systems require a combination of different tools^52^, such as MS-cleavable cross-linkers^53, 54^, enrichment strategies of crosslinking products, and dedicated software^44, 56^. System-wide XLMS studies have now become popular for a broader community due to the rapid technological advances in MS instrumentation, commercially available MS-cleavable crosslinkers, such as disuccinimidyl dibutyric urea (DSBU or BuUrBu)^56^ or disuccinimidyl sulfoxide (DSSO)^57^.

The complexity of MS/MS spectra of cross-linked peptides14 together with the large number of theoretical reaction pose great challenges for the software. Even for isolated proteins or small complexes, the correct false-discovery rate (FDR) estimation is not easily achieved^58, 59^ and a manual validation of spectra might be beneficial. XLMS experiments on complex protein mixtures usually yield thousands of cross-links, making the inspection of individual spectra impossible. Recently DSBU and MeroX have been combined for developing XLMS workflows to study isolated proteins or protein complexes.^19,60-62^

Our in-house-software MeroX (www.StavroX.com) was especially designed for the analysis of MS/MS-cleavable crosslinkers. With previous versions of MeroX, analyses were limited to 500-1000 proteins in the sequence database. Therefore, we aimed at upgrading MeroX to perform truly automated and reliable cross-link analysis at a proteome-wide scale. MeroX 2.0 relies on two novel algorithms dedicated to MS-cleavable cross-linkers, termed RISEUP and proteome-wide mode, and is fully scalable from single proteins to systemwide XLMS studies. The new algorithms also comprise a series of novel filtering steps. MeroX 2.0 analyzes highresolution full scan mass spectra for improved monoisotopic peak picking and re-associates the corrected m/z values of precursor ions to the relevant MS/MS spectra. Implementing this precursor mass correction yields ∼ 20% more identification of crosslink spectra, while reducing false positives due to incorrect mass-to-charge ratios of molecular ions. After having identified the correct monoisotopic signals, MeroX 2.0 permits a mass recalibration at MS and MS/MS levels. As the distribution of m/z deviations is rarely centered at zero, a linear recalibration shift allows narrowing the m/z tolerance windows imposed. This reduces the decoy matches and the overall computational time requirements, while increasing the number of cross-link identifications and amplifying their reliability. MeroX 2.0 automatically draws detailed, interactive protein network maps, thereby facilitating deep cross-link analyses. MeroX 2.0 also incorporates the well-established algorithm of StavroX^63^ enabling the identification of cross-links from classical, non-MS-cleavable reagents, such as bis(sulfosuccinimidyl)suberate (BS3) and disuccinimidyl suberate (DSS). At the same time, the possibility for fine-tuning specific search parameters as well as the interactive visualization of fully assigned MS/MS spectra have been improved.

We evolved our previous XLMS protocol^19^ for investigating isolated proteins and protein complexes to establish an easyto-use protocol for proteome-wide XLMS analysis based on the MS-cleavable, commercially available DSBU cross-linker, size exclusion chromatography (SEC) for cross-linking product fractionation, and a nano-HPLC/nano-ESI-Q-Exactive Plus Orbitrap mass spectrometer (Thermo Fisher Scientific) using MeroX 2.0 (Figure 1). We optimized the workflow by reducing the sample handling operations between all protocol steps. Most importantly, our protocol does not require any higher order fragmentation schemes but relies exclusively on MS/MS experiments. This results in a very fast, easy-to-handle workflow that is broadly applicable to many laboratories worldwide having access to mass spectrometers with tandem MS capabilities.

**Figure 1.**
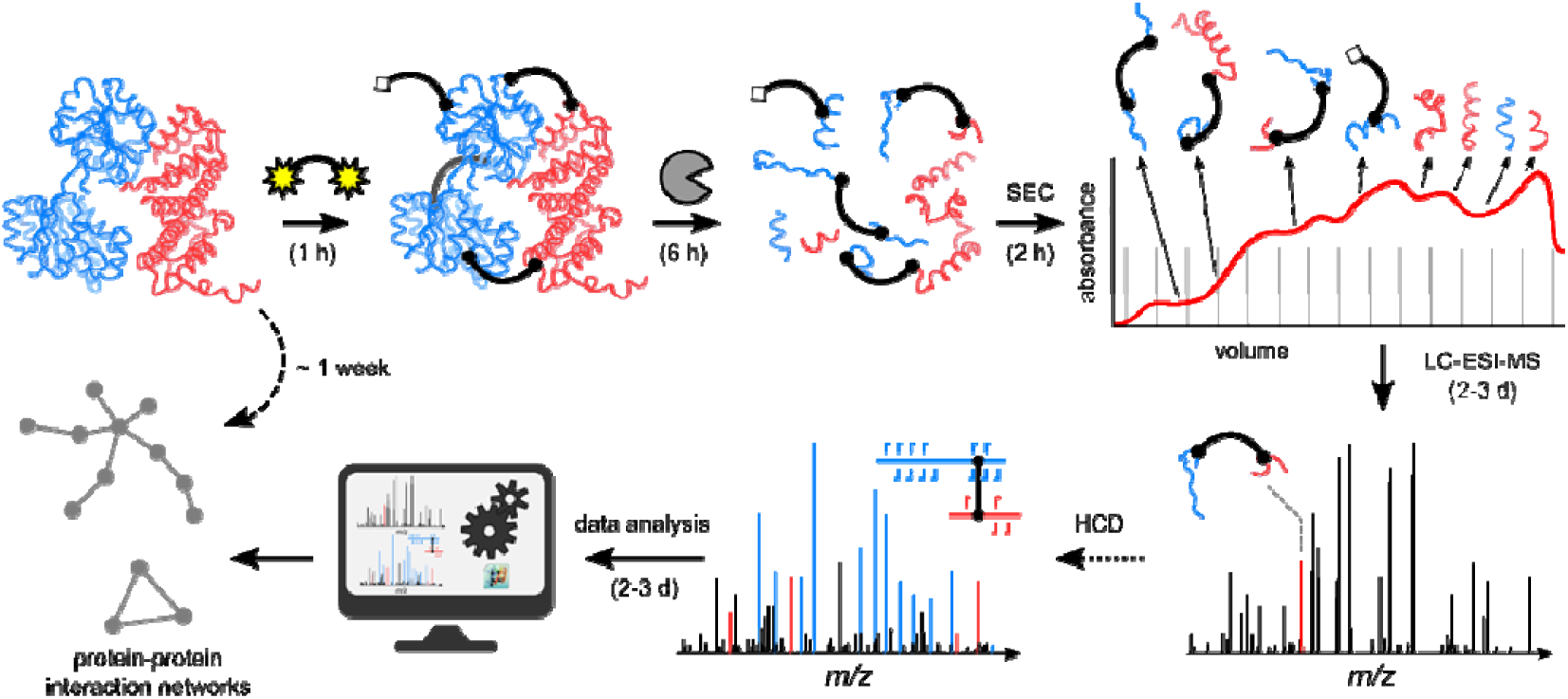
General workflow of proteome-wide cross-linking/MS studies using MS/MS-cleavable linkers and MeroX 2.0.

Our protocol has been applied to study protein complexes in extracts of *Drosophila* melanogaster embryos. Early *Drosophila* embryos (0-2 h) are transcriptionally silent. In that they solely depend on maternal RNA and proteins provided during oogenesis. All processes until transcriptional genome activation can therefore only be regulated at a post-transcriptional level e.g. RNA stability control or translational regulation. Extracts used in this study are active in translation of mRNAs and reconstitute translational regulation of the nanos-mRNA that is required for posterior patterning^64, 65^. Protein-protein interactions (PPIs) identified in this system might aid in understanding the complex nature of regulation during early embryonic development.

In this complex system, we identified 5,129 unique crosslinking sites comprising 694 protein-protein-interactions at 5% FDR. Interestingly 492 interactions are novel interactions occurring during early embryo development before tran criptional genome activation.

## RESULTS AND DISCUSSION

### Proteome-Wide Cross-link Analysis of Drosophila Extracts

To explore the potential of our experimental workflow using the MeroX 2.0 software on a proteome-wide level, we used extracts of *Drosophila* melanogaster embryos at an age of 0-2 h. In these extracts, several biochemical processes can be successfully recapitulated64. These processes include translation from in-vitro-synthesized mRNA, translational repression, and sequence-specific deadenylation. Translation is a highly demanding task requiring an immense number of correct protein-protein interactions. We confirmed that the extracts used for the cross-linking reactions are active in translation as well as a sequence-specific translational repression (Figure 2A). For the cross-linking reactions we employed identical conditions as for translational repression assays. Cross-linking was performed at a total protein concentration of ∼ 20 g/l and 1 mM of DSBU (Material and Methods), corresponding to a ∼ 2.5-molar excess of DSBU over protein. We were able to reproducibly observe the formation of higher-order protein complexes without over-cross-linking the samples as judged by SDS-PAGE analysis (Figure 2B).

**Figure 2.**
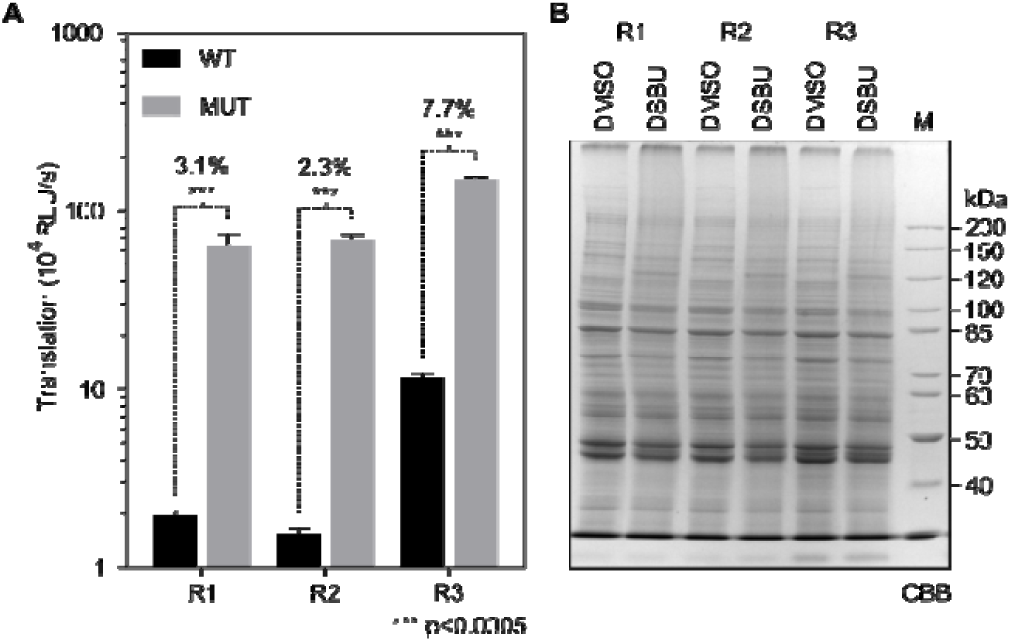
(A) Three biological replicates of *Drosophila* melanogaster embryo extract were tested for translation and smaug-dependent translational repression. 2 nM of either wildtype RNA (WT translationally repressed) or mutated RNA (MUT containing two point mutations in the regulatory sequence) were incubated in extracts and translation was started by completion of an ATP-regenerating system. Translational repression indicated above each pair of data is remaining translation in WT (TLWT/TLMUT) and ranges from 2.3% to 7.7%. (B) Crosslinking of the three replicate extracts was performed at ∼20 g/l protein and 1 mM of DSBU or equivalent volumes of DMSO as control. Cross-linked samples were separated by SDS-PAGE and stained with Coomassie Brilliant Blue.

### Cross-linking Workflow

We optimized our existing crosslinking workflow that had originally been developed for analyzing purified proteins and protein complexes^19^ in respect to a fast and reliable proteome-wide analysis of cross-links (Figure 1). For an in-depth analysis of complex samples, such as whole cell lysates, separation and/or enrichment steps of cross-linked peptide species are essential. For the enrichment of cross-linked peptide, we applied size-exclusion chromatography (SEC)^60, 66, 67^. Without any further processing, samples were subjected to high-resolution tandem mass spectrometry (nano-HPLC/nano-ESI-MS/MS). Raw data were converted to standard mzML format and analyzed using the updated MeroX 2.0 version. This new version is capable of identifying crosslinks from highly complex samples with an entire proteome. From cross-linked *Drosophila* embryo extracts, we detected 18,037 cross-link spectrum matches (XSMs) that can be reduced to 5,129 unique cross-linked residues in biological triplicate experiments at 5% FDR (Figure 3A and B, Table 1). Among these, 1,237 inter-protein cross-linking sites were identified that contain valuable information on PPIs. The remaining 3,892 intra-protein cross-links deliver information on the conformation of proteins in their cellular environment. It is important to point out that these numbers exclude crosslinks that are composed of consecutive amino acid sequences, which are usually reported in other studies. In total, those account for 35% more XSMs (additional 6,493 XSMs) and 26% more unique cross-linked residues (additional 1,343 unique sites).

**Table 1.** Summary of unique cross-linking sites, crosslinked proteins, and PPIs, identified by MeroX 2.0. The numbers are shown separately at different FDR and for different database sizes. ID: identified proteins; DP: *Drosophila* proteome; CP: consecutive peptide.

| FDR cutoff | Database size | Cross-linked peptides, spectral matches | Total unique cross-linking sites (excluding CP) | Intra-molecular unique cross-linking sites (excluding CP) | Inter-molecular unique cross-linking sites | PPIs |
| --- | --- | --- | --- | --- | --- | --- |
| 5% | ID (9,536 proteins) | 18,037 | 5,129 | 3,892 | 1,237 | 692 |
| 5% | DP (21,973 proteins) | 16,184 | 4,776 | 3,615 | 1,161 | 664 |
| 1% | ID (9,536 proteins) | 9,945 | 3,098 | 2,568 | 530 | 255 |
| 1% | DP (21,973 proteins) | 8,636 | 2,845 | 2,358 | 487 | 238 |

**Figure 3.**
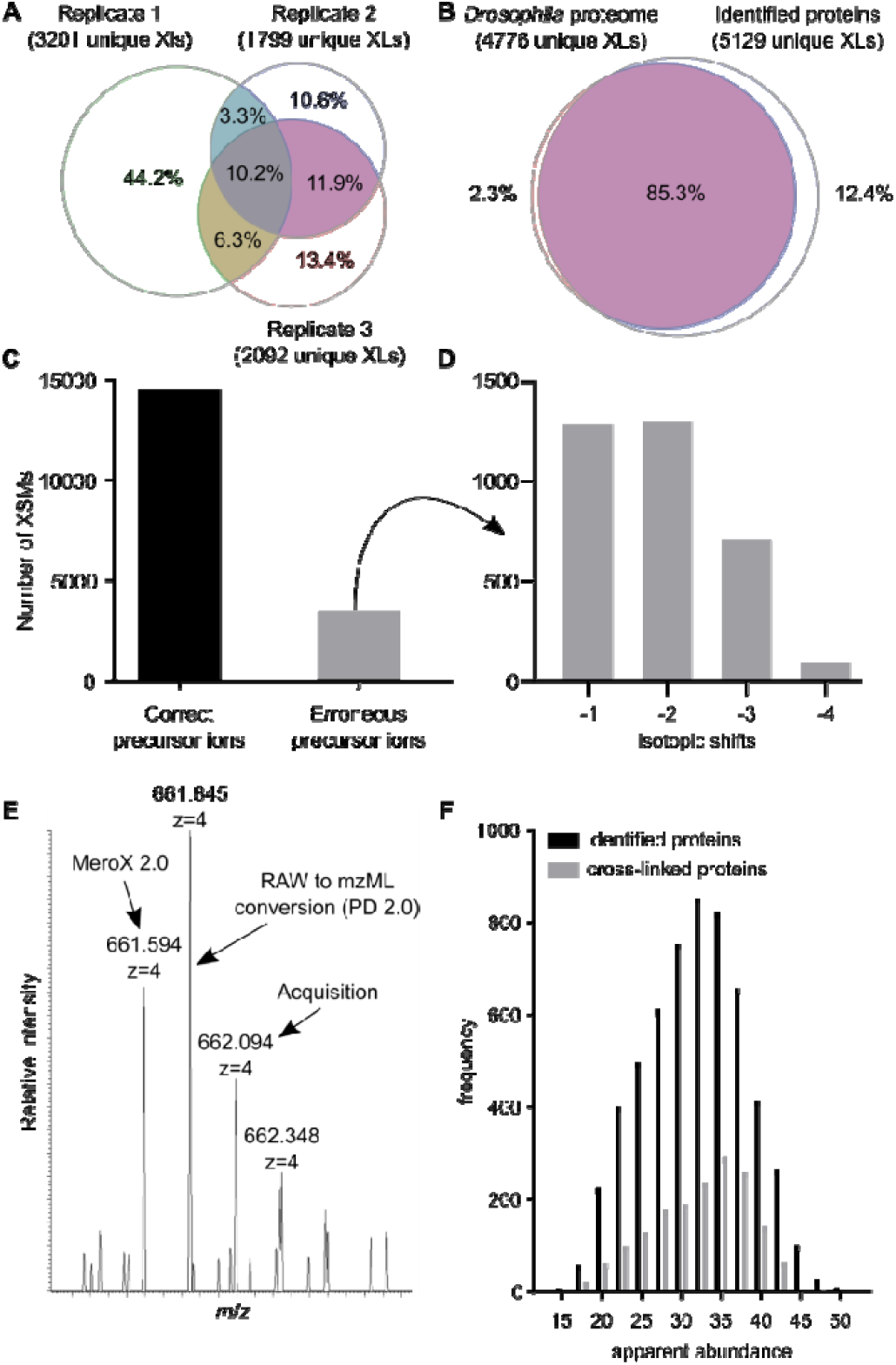
(A) Cross-linking analyses were performed in triplicates; the overlap of unique cross-linking sites between replicates is shown. (B) Comparison of data analysis based on the set of identified proteins (5,984 protein entries) or the complete *Drosophila* proteome (21,973 protein entries). (C) Identification of cross-linked peptides in spectra with inaccurately assigned precursor signals. (D) Number of isotopes, with the algorithm correcting precursor masses. (E) Representative mass spectrum where the precursor mass was corrected by MeroX. (F) Abundance distribution of proteins identified in *Drosophila* and proteins found in cross-links. The distribution of protein intensities in cross-links are skewed towards higher intensities.

Cross-linked amino acids that are separated by only a few residues usually significantly increase the number of overall cross-link identifications, but delivers low-impact geometrical constraints.^14^ MeroX 2.0 offers the unique option to exclude cross-links that are composed of consecutive amino acid sequences, to generate lists of identified cross-links that focus on the structurally most relevant species.

### Evaluation of Cross-linking Sites

To expedite computational analysis we restricted MeroX 2.0 analyses to a database with a reduced size containing 9,536 protein entries that had been identified in the cross-linked samples by MaxQuant. Comparing the outcome to the same analysis against the whole *Drosophila* proteome (21,973 protein entries), we observed an overlap of 85% (Figure 3B). We identified 4,776 unique crosslinks using the *Drosophila* proteome versus 5,129 unique cross-links with the set of identified proteins (Table 1). We therefore conclude that the lower number of cross-links in the whole proteome search is caused by an accumulation of decoy and false-positive identifications that were filtered at the FDR level, while discarding true cross-links. The same analysis at 1% FDR yielded 3,098 unique cross-linked residues. Data throughout the paper discuss the cross-links obtained at 5% FDR using abovementioned database containing 9,536 proteins sequence entries (including isoforms).

The limited overlap of cross-links we identified in three biological replicates, i.e. three different *Drosophila* embryo extracts (Figure 3A), reflects the variability in embryo treatment, extract production, complexity of the cross-linked peptide mixture on a system-wide scale, and the mild conditions used in our cross-linking reaction.

We aimed at using a low molar excess of DSBU in order to limit an eventual perturbation of native protein conformations and PPIs. This is in agreement with a previous report describing a cross-linker concentration of 1 mM to be optimally suited for analyzing human cell lysates.^20^

On the other hand, low cross-linker concentrations prevent targeting all lysine residues in a single experiment and limit the overlap between biological replicates.

SEC fractionation dramatically increases the number of identified cross-links in system-wide XLMS studies. Intriguingly, preliminary analyses of unfractionated cross-linked *Drosophila* embryo extract yielded only a low number of cross-links (data not shown). SEC does not completely resolve the issue of sample complexity as we could demonstrate by repeating LC/MS/MS analyses of two selected SEC fractions. MeroX 2.0 revealed that the overlap of cross-linked residue combinations in these technical replicates is ∼ 45% (Figure S1). Employing Orbitrap Q-Exactive HF-X or Orbitrap Fusion Lumos instruments may substantially increase the number of crosslink identifications and the number of PPIs accessible by a XLMS experiment. We used the MaxQuant software to quantify the proteins present in the sample.^68^ Importantly, abundant proteins are enriched among the observed cross-links, but identifications are not limited to high-abundant protein species (Figure 3F). Nevertheless, we identified cross-links including also low abundant proteins.

### Mass Correction

Precursor ions of cross-linked peptides are usually low-abundant in mass spectra compared to those of linear peptides. Monoisotopic peak picking might fail on low abundant ions during conversion of MS data from RAW files.^69, 70^ In 20% of all spectra representing cross-linked peptides, the precursor mass had to be corrected, mostly by one to three isotope peak shifts (Figure 3C-D). This percentage is consistent with previously reported rates of precursor ion misassignment of low abundant species.^69, 70^ This is exemplified in Figure 3E: The 4+ charged ion at *m/z* 661.594 was not detected as the monoisotopic signal during conversion using Proteome Discoverer 2.0. Strikingly, MeroX identified the correct monoisotopic signal and adjusted the reported precursor mass. The corresponding MS/MS spectrum could then be assigned to the intra-protein cross-link between K173 and K182 of the Drosophila elongation factor eIF3g1 (Figure 3D, Figure S2 and S3). Consequently, the mass correction algorithm implemented in MeroX 2.0 increases the number of cross-link identifications by avoiding false positives due to a wrong precursor mass selection.

### Cross-link Validation for the Drosophila Ribosome

We evaluated the cross-links involving ribosomal proteins by mapping them into the high-resolution structure of the *Drosophila* 80S ribosome^71^ as an example of a large protein assembly (Figure 4). We were able to map 232 (167 intra- and 65 inter-molecular) unique cross-linked residues in the struct rally resolved portion of the *Drosophila* 80S ribosome revealing that 215 cross-links (93%) are compatible with the known high-resolution structure (Figure 4D). The distance dis ribution obtained from the cross-linking data significantly differs from a random distribution of interand intra-molecular crosslinks within the ribosome (Figure 4D). Interestingly, 15 of the remaining 16 cross-links exceeding the maximum Cα-Cα distance of 32 Å are not randomly distributed over the 80S ribosome. They cluster in the regions of L10Ab/L11/L36A and L6/L35a/L17/L28 ribosomal proteins, and the eukaryotic elongation factor eEF2. In each of these three regions the overlength cross-links possess identical orientations, being almost parallel to each other (Figure 4B). This indicates that most likely they do not represent artificial cross-links, but rather suggest slightly different conformations in these protein regions. Based on five inter-molecular cross-links ((K557(eEF2)-K80(S23), K484(eEF2)-K40(L12), K498(eEF2)-K74(L23), K498(eEF2)-K75(L23), and K498(eEF2)-K113(L23)), we could also confirm the topology of the eEF2–80S complexes^71^. The GTP-binding elongation factor eEF2 mediates the translocation of peptidyl-tRNA and is the target of the Diphtheria toxin^72^. Our cross-links correctly locate this essential factor at the edge between the small and large subunit of the 80S ribosome of *Drosophila*. Six crosslinks identified in eEF2 involve residues of the C-terminal domain and they systematically exceed the maximum C_α_-C_α_ distance of 32 Å for the DSBU cross-linker (Figure 4C). As this suggests the existence of a different conformation of eEF2 in solution, we mapped these cross-links into the high-resolution structure of eEF2 from Saccharomyces cerevisiae (pdb entry 1zm2).

**Figure 4.**
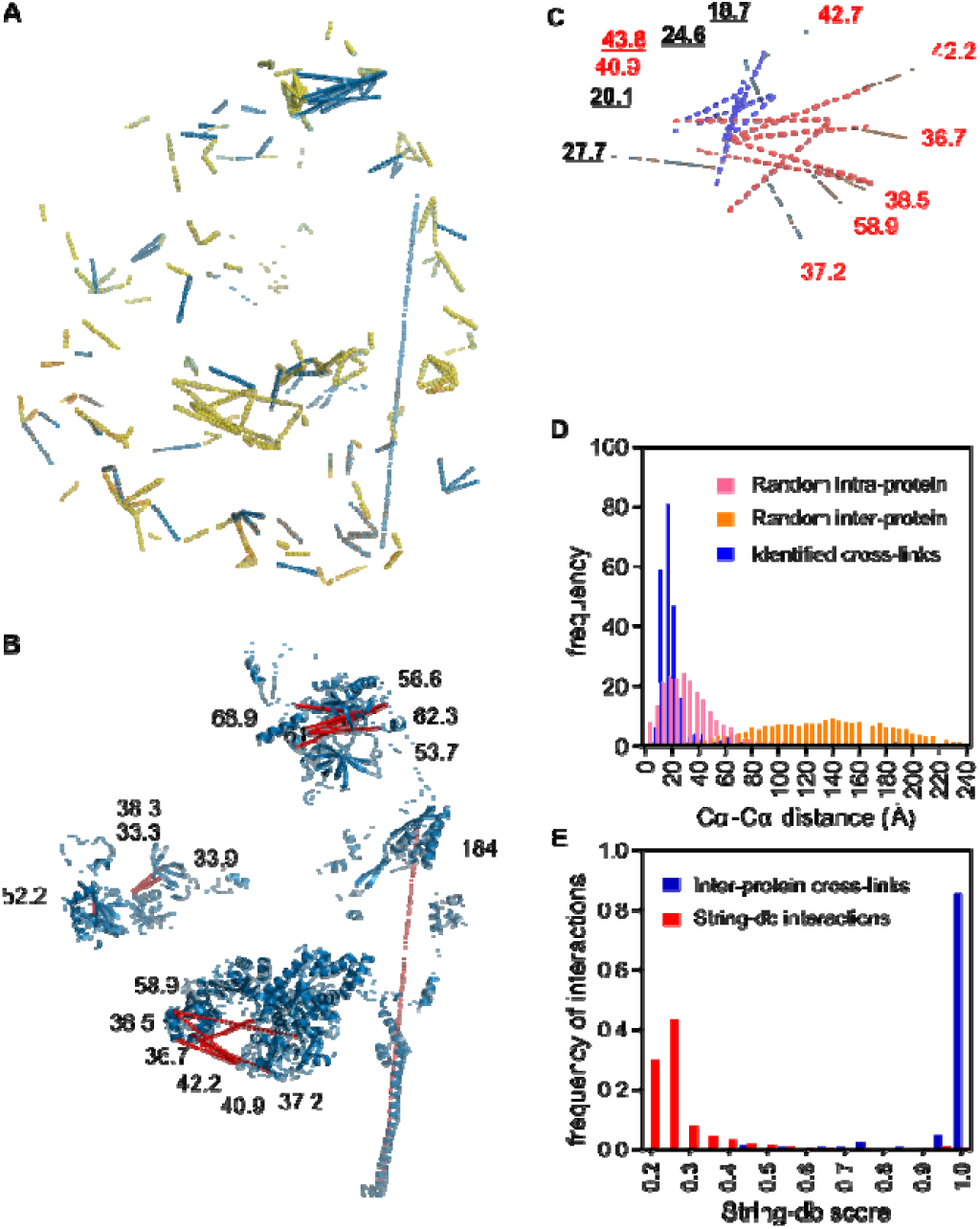
(A) All cross-links found in the ribosome were mapped onto the cryo-EM structure of the *Drosophila* ribosome (PDBcode: 4v6w). 65 inter-protein cross-links are shown in blue, 167 intra-protein cross-links in yellow. (B) In total, 16 cross-links exceeded the limit of 32 Å (shown in red). The subunits involved in these over-length cross-links are highlighted. (C) Six crosslinks with a length above 30 Å were detected within the structure of eEF2 (orange). In the structure of the yeast eEF2 (PDB entry: 1zm2), four of those cross-links are below the distance threshold (dashed blue lines, distances in the Yeast eEF2 structure are shown in bold and underlined). (D) C_α_-C_α_-distance distribution of cross-links identified in comparison with random distances of intra- and inter-protein cross-links. (E) All inter-protein crosslinks were compared to the STRING database containing known PPIs. 202 of 694 unique PPIs were also present in the STRING database. The distribution of STRING database scores was plotted for all *Drosophila* proteins as well as for the subset of identified cross-links. PPIs with high confidence were found to be clearly enriched among the cross-linked protein pairs identified.

In fact, ∼ 70% of the eEF2 amino acids sequence of *Drosophila* melanogaster is identical to that of Saccharomyces cerevisiae and they share a highly similar fold^71^ (Figure 4C). The eEF2 N-terminal domain of *Drosophila* melanogaster and Saccharomyces cerevisiae that binds to the nucleotide is largely superimposable, while the eEF2 C-terminal domain of Saccharomyces cerevisiae shows a slightly different orientation (Figure 4C). This results in a more compact structure of Saccharomyces eEF2 compared to that of *Drosophila* eEF2. Interestingly, four of the overlength intramolecular cross-link agree with the structure of the yeast ortholog (Figure 4C). This suggests the existence of a more compact conformation of *Drosophila* eEF2 in solution, similar to the conformation of Saccharomyces eEF2.

Cryo-EM data had suggested that the Vig2 protein binds the 80S ribosome of *Drosophila* on the head of the 40S subunit mRNA in proximity to the mRNA channel.^71^ This discloses a novel metazoan function of translation regulation of this protein.^71^ We could confirm the location of Vig2 on the head of the 40S subunit by identifying seven inter-protein cross-links (K62(Vig2)-K202(S3A), K297(Vig2)-K203(S3), K302(Vig2)K29(S19a), K392(Vig2)-K101(S20), K111(Vig2)-K66(S26), K111(Vig2)-K12(S28b), and K111(Vig2)-K27(S28b)) between Vig2 and ribosomal proteins.

On top of that, we detected 761 intra- and 559 inter-molecular cross-links involving flexible and dynamic peripheral ribosomal proteins and additional ribosome interacting partners, for which the structures have not been solved yet. All captured protein interactors of the 80S ribosome in *Drosophila* embryos are shown in Figure S4. It has been reported that orthologs of Rpl23a (L23A), along with other surface-exposed components at the peptide exit tunnel, serve as an evolutionary conserved docking site for many bacterial and eukaryotic chaperones, processing enzymes, and targeting factors that bind to nascent chains on translating ribosomes^73^. Interestingly, L22 and L23A are located directly at the peptide exit tunnel, so two explanations are conceivable for the high amount of cross-links involving L22 and L23A. The cross-linked proteins are histones or modifying enzymes that are needed cotranslationally, or alternatively, nascent chains were cross-linked to the ribosome in our experiments. When checking the proteins that were cross-linked to L23A (Figure S4) we neither identified chaperones, nor any modifying enzymes, so we hypothesize that our cross-linking experiments might have captured a snapshot of the ribosome during translation.

### Cross-link Validation for Protein Oligomers

Mapping crosslinks in high-resolution structures of protein complexes comprised of two or more identical proteins is challenging. In a system-wide XLMS experiment, it is not possible to distinguish between intra- and inter-molecular cross-links for homooligomers. To exemplify this often-overlooked limitation of the XLMS technique, we tentatively mapped cross-links in the structural model of aldolase (pdb entry 2pyo). *Drosophila* aldolase is a homo-tetramer of 158 kDa catalyzing the aldol cleavage of fructose-l,6-bisphosphate.74 We identified a total of nine cross-links in aldolase of *Drosophila* embryos. Two of them, composed of identical amino acid sequences, can be unambiguously assigned to inter-protein cross-links with excellent agreement with the aldolase 3D-structure (Figure 5A), while the remaining seven cross-links were assigned both as intra- and inter-protein cross-links. We measured the Cα-Cα distances of these ambiguous cross-links within a monomer of *Drosophila* aldolase and between the two of them in the tetramer. The Cα-Cα distances measured as intra-protein crosslinks range between 9.6 and 19.0 Å, while all Cα-Cα distances measured as inter-protein cross-links largely exceed the limit of 32 Å of DSBU (Figure S5B). This suggests that these crosslinks involve amino acid pairs located in the same aldolase chain.

### PPIs in Drosophila embryos

We compared the identified inter-protein cross-links to known PPIs in the String database version 10 (STRING-db.org). The 1,237 unique cross-linking sites identified herein represent 694 unique PPIs and 202 of these interactions are also found in the String database The STRING database scores of these 202 cross-links are among the highest scores reported in this PPI database (Figure 4E). All scores range above 0.4 (String medium confidence), while 90% of all observed protein pairs have a STRING database score higher than 0.9 and are therefore highly confident interactors. Strikingly, 492 interactions are not present in the STRING database, suggesting that these interactions present novel protein-protein interactions in the *Drosophila* embryo. The complete list of PPIs identified by this work is provided in Table S1 After the validation of our cross-links on the highresolution structure of 80S ribosome, the high confidence of the identified PPIs confirms the reliability of our XLMS approach. Thus, the unprecedented 492 PPIs captured in this work can be mined while studying the functional protein organization and topology in early embryogenesis.

## CONCLUSIONS

We present a proteome-wide XLMS workflow that can easily be performed within the timeframe of one week and is based on commercially available reagents and equipment. We upgraded the freeware tool MeroX 2.0 to conduct fully automated, fast, and reliable large-scale protein-protein cross-linking analyses. We applied this integrated workflow to capture PPIs at a system-wide level of early *Drosophila* embryos undergoing maternal-to-zygotic transition. In total, we identify 5,129 unique cross-linked residues involving proteins in a wide dynamic range of the cellular proteome. Our results were validated based on the high-resolution structure of 80S ribosome and the STRING interaction database. 93% of all crosslinks fall within the maximum distance range for the DSBU linker and the remaining 7% give insights into local conformational changes of ribosomal components in solution. More than 90% of the PPIs that are detected by our work and are also present in the STRING database are among the top 1% entries in the STRING interaction database with the highest confidence score (> 0.9). Conclusively, we unveiled 492 unprecedented PPIs defining the topology of novel protein complexes in early *Drosophila* embryogenesis.

## MATERIALS AND METHODS

### Cross-linking

*Drosophila* embryo extracts were prepared as previously described^65^. Extracts (40%) were used in crosslinking reactions with 26 mM HEPES, pH 7.4, 81 mM potassium acetate, 1.6 mM magnesium acetate, 1.3 mM ATP. DSBU (stock of 50 mM in DMSO) was added to a final concentration of 1 mM and incubated for 60 minutes at room temperature. Cross linked samples were flash-frozen and stored at −80°C.

### Digestion and Fractionation

150 µl of cross-linked sample (∼3 mg of total protein) were digested by SMART Digest Trypsin Kit (Thermo Fisher Scientific), by adding 450 µL of SMART Digest buffer containing 50 µg of beads at 70°C for 4 h. The sample was cooled to room temperature and trypsin beads were removed by quick centrifugation. For reduction of cysteines, 4 mM of DTT was added and incubated for 30 min at 56°C. 8 mM iodoacetamide was added and incubated 20 min at room temperature in the dark. To prevent overalkylation, 4 mM of DTT were added. 600 µL of digested cross-linked sample were separated on an ÄKTA Pure system (GE Healthcare) using a Superdex 30 Increase 10/300 GL size exclusion chromatography column. The column was equilibrated and operated with 3% (v/v) acetonitrile, 0.1% (v/v) TFA. 18 fractions (500 µL each) were collected from 8 to 20 mL into 96 deep-well plates.

### Mass Spectrometry

SEC fractions were analyzed by LC/MS/MS on an UltiMate 3000 RSLC nano-HPLC system (Thermo Fisher Scientific) coupled to an Orbitrap Q-Exactive Plus mass spectrometer (Thermo Fisher Scientific) equipped with Nanospray Flex ion source (Thermo Fisher Scientific). Peptides were separated on reversed phase C18 columns (precolumn: Acclaim PepMap 100, 300 μm × 5 mm, 5μm, 100 Å (Thermo Fisher Scientific); separation column: packed Picofrit nanospray C18 column, 75 μM × 250 mm, 1.8 μm, 80 Å, tip ID 10 µm (New Objective)). After washing the precolumn for 30 minutes with water containing 0.1 % (v/v) TFA at a flow rate 50 μl/min and a pre-column temperature 50°C, peptides were eluted and separated using a linear gradient from 3% to 42% B (with solvent B: 0.1% (v/v) formic acid and 85% (v/v) acetonitrile) with a constant flow rate of 300 nl/min over 360 min, 42% to 99% B (5 min) and 99% B (5 min). The separation column is kept at 45°C using an external column heater (Phoenix S&T).

Data were acquired in data-dependent MS/MS mode with stepped higher-energy collision-induced dissociation (HCD) and normalized collision energies of 27%, 30%, and 33%. Each high-resolution full scan (m/z 299 to 1799, R = 140,000 at m/z 200) in the orbitrap was followed by 10 high-resolution product ion scans (R = 35,000), starting with the most intense signal in the full-scan mass spectrum (isolation window 2 Th); the target value of the automated gain control was set to 3,000,000 (MS) and 250,000 (MS/MS) and maximum accumulation times were set to 100 ms (MS) and 250 ms (MS/MS). Precursor ions with charge states <3+ and >7+ or <3+ and >5+ were excluded from fragmentation of SEC fractions 1-3 and 4-18, respectively. Dynamic exclusion was enabled (duration 60 seconds, window 2 ppm).

### Data Analysis

All proteins present in the samples were identified using the MaxQuant software^68^ with the *Drosophila* proteome as database (uniprot.org). Based on this analysis, a fasta file containing all identified proteins, including isoforms was downloaded from uniprot.org. For cross-linking analysis, mass spectrometric *.raw files were converted to mzML using Proteome Discoverer 2.0. MeroX analysis was performed with the following settings: Proteolytic cleavage: C-terminal at Lys and Arg with 3 missed cleavages, peptides’ length: 5 to 30, static modification: alkylation of Cys by IAA, variable modification: oxidation of M, cross-linker: DSBU with specificity towards Lys and N-termini, consecutive sequences were discarded, precursor mass accuracy: 4 ppm, product ion mass accuracy: 8 ppm (performing mass recalibration, average of deviations), signal-to-noise ratio: 1.5, precursor mass correction activated, prescore cut-off at 55% intensity, FDR cut-off: 1%, 5%, and minimum score cut-off: 70.

## ASSOCIATED CONTENT

### Supporting Information

The Supporting Information is available. It contains the description of the software MeroX 2.0, Table S1 with all identified protein-protein interactions, as well as Figures S1-S7, showing a comparison of technical replicates of SEC fractions, details on precursor mass correction, the ribosomal interaction network of *Drosophila*, mapping of cross-links in the crystal structure of the aldolase homotetramer, filtering steps applied by MeroX and properties of the MS-cleavable cross-linker DSBU.

Raw data are deposited at PRIDE.

## AUTHOR INFORMATION

### Author Contributions

The manuscript was written through contributions of all authors. All authors have given approval to the final version of the manuscript.

## ACKNOWLEDGMENT

A.S. acknowledges financial support by the DFG (project Si 867/15-2). C.I. is funded by the Alexander von Humboldt Foundation. M.G. was supported by the RTG 1591 at the Martin Luther University Halle-Wittenberg. The authors are indebted to Prof. Elmar Wahle for continuous support.

---

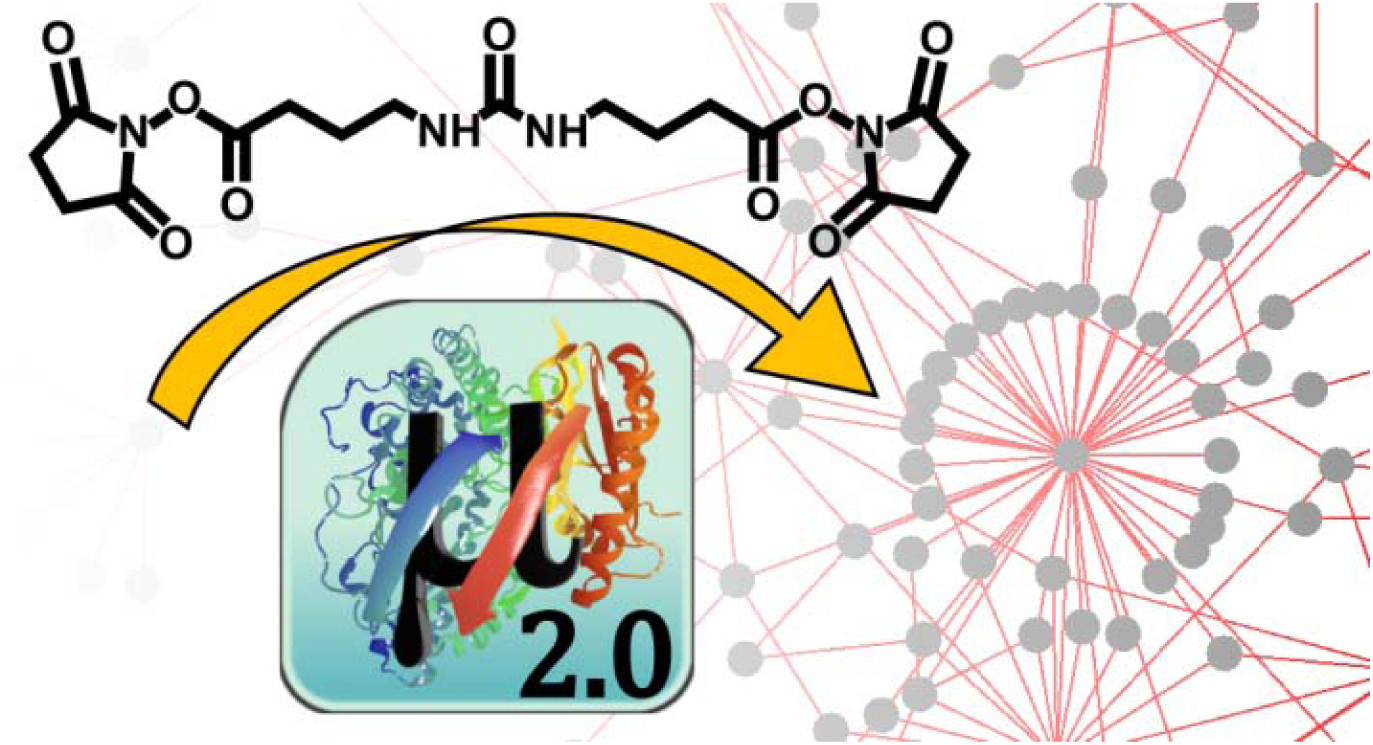
Graphical Abstract.

