## Supplementary Information for "A Simple Cross-Linking/Mass Spectrometry Workflow to Study System-Wide Protein Interactions"

### Software Description

The MeroX software[61] was originally designed for purified proteins and small protein complexes and was therefore less well suited for a proteome-wide identification of cross-links. To advance MeroX for performing proteome-wide analyses, several important features had to be included.

*Novel Modes.* -- Two novel modes were implemented, termed “RISEUP mode” and “proteome-wide mode” to complement the already existing modes of data analysis (quadratic mode and RISE mode[61]). The RISEUP mode is an advanced RISE mode that is able to infer peptide masses from incomplete reporter ion patterns. The proteome-wide mode relies on the presence of reporter ion pairs for at least one of the two peptides. Subsequent filtering steps allow the analysis of much larger databases, such as the human proteome, than previously attainable with MeroX (**Fig. S6**). The different modes form a hierarchic system, allowing a specific adjustment for each protein system under investigation: The quadratic mode performs best for up to 10 proteins as it does not rely on the presence of reporter ions in the fragment ion mass spectra. Thus, the quadratic mode is the algorithm of choice for identifying non-MS-cleavable cross-links. The RISE and RISEUP modes are optimally suited for up to ~500 proteins, while the proteome-wide mode allows identifying cross-links from several thousands of proteins in a database.

*In-silico Proteolysis.* -- One of the key steps that had to be optimized in the new MeroX version was *in-silico* proteolysis. For a fast analysis, all cross-linkable peptides are pre-calculated by an *in-silico* digestion and stored in the memory. We implemented a peptide database that contains all necessary information on all peptides with minimal redundancies. These databases can be generated at once, stored locally, and used every time the same organism and protease combination are used. For tryptic digestion (up to three missed cleavages) of the *Drosophila* proteome, 3.8 million peptides are generated from 21,973 proteins within 5 minutes.

*Preprocessing.* -- All spectra are preprocessed before data analysis. After noise reduction, charge states of the fragment ions are determined and only monoisotopic signals are retained[19]. Discrepancies might originate from the different extraction procedures of MS data from RAW data using different software tools.[62, 63] Conversion software might fail in selecting monoisotopic peaks in the mass spectra, which is of particular relevance for low abundant precursor ions, such as cross-linked peptides. To correct the eventually false assignments of precursor ions, MeroX analyzes the corresponding mass spectra that are exported to mzML or mzXML file formats by searching for the correct monoisotopic signals (**Fig. 3, Fig. S2**). For other input formats containing exclusively MS/MS spectra, e.g. mgf, MeroX iterates through potential precursor to identify a matching cross-linked peptide pair.

*Reduction of Cross-link Candidates.* -- Reducing the search space is undoubtedly one of the most pivotal requirements for conducting fast proteome-wide analyses on standard PCs. MeroX uses several filters to reduce the number of cross-link candidates that will be scored afterwards (**Fig. S1**). First, each spectrum is scanned for reporter ion pairs originating from MS-cleavable cross-linkers that match the mass difference between two fragments of the same peptide. For the DSBU cross-linker, this mass difference corresponds to 25.979 u (**Figure S7**). Only ions with a defined charge state are utilized for this search. From each reporter ion pair, the corresponding peptide mass can be inferred and all peptides in the database matching the identified fragment ion pattern are considered for further analysis. In an initial search round, the peptides identified are compared to the spectrum in an open mass modification search. The resulting peptide score $S_{peptide}$ is calculated as follows:

$$S_{peptide} = 10 \cdot coverage \cdot \sum_{i}^{n} intensity_i$$

where *coverage* corresponds to the fraction of cleaved peptide bonds and $intensity_i$ is the relative intensity of signals identified in an MS/MS spectrum. All peptides below a certain threshold (with a default value of 10) will not be considered further. For the filtered peptides, the mass of the second peptide is inferred by subtracting the mass of the first peptide from that of the precursor ion. Therefore MeroX 2.0 enables an efficient identification of cross-links even from MS/MS spectra where only one doublet or reporter ion is visible (**Figure S6**). All peptides in the database matching the mass of the second peptide are also filtered by the peptide score with the same threshold as before. All remaining pairs of peptides are sorted based on increasing score.

*Prescoring.* As an additional filter, the prescoring algorithm for MeroX was modified to meet the needs of the novel RISEUP and proteome-wide modes (see above). Signals in the precursor ion isolation window (*m/z* ± 4 Th) are removed from the spectrum. This avoids the interference of coisolated species, which often remain intact under the collisional activation conditions used for peptide fragmentation. To calculate the prescore, all signals above 10% relative signal intensity of the remaining base peak are considered for prescoring. The intensities of all identified signals in one spectrum are added and divided by the total ion current. Only spectra with a minimum percentage (55% default value) of the total ion current assigned to fragment ions are considered for subsequent scoring. Finally, MeroX 2.0 scores the filtered cross-linking candidates by a fragment ion intensity-based algorithm as described previously (Iacobucci et al., *Nature Protocols,* 2018; doi: 10.1038/s41596-018-0068-8).

Also, different features of the annotated MS/MS spectrum are quantified, i.e., the number of identified signals, ion series, reporter ion presence and intensity. The analysis is repeated on spectra where all *m/z* values have been shifted by a constant value to account for random overlaps. The results for the original spectra and mass shifted spectra are compared and used in a weighted scoring function to obtain the final score. A score value of ~ 100 represents a good match.

*False-Discovery Rate.* -- The estimation of false-discovery rates (FDR) is based on a target decoy approach. The complete protein database is used to generate decoy protein entries by scrambling the amino acid sequence. By default, MeroX scrambles the amino acid sequences of each protein, while keeping the cleavage sites in place. In cross-link analyses, decoy candidates comprise decoy-decoy candidates as well as mixed target-decoy species. The number of decoy candidates is therefore three times higher than the number of target candidates. Consequently, the FDRs for a set of candidates above a certain score cut-off are calculated by counting all decoy and target candidates above the threshold using the following equation:

$$FDR = \frac{N_{decoy}}{3 \cdot N_{target}}$$

For the proteome-wide mode, FDR values change in dependence of the minimum peptide score applied. MeroX 2.0 iterates through different minimum peptide score cut-offs (up to 80) and calculates the number of cross-link candidates above the FDR cut-off applied (1% or 5% FDR). The minimum peptide score cut-off yielding the highest number of cross-link candidates at the defined FDR is finally applied and the results are filtered by a defined score cut-off.

*Precursor Mass Correction.* -- Precursor ions of cross-linked peptides are usually of low intensity compared to these of linear peptides. Monoisotopic peak picking might fail on low abundant ions during conversion of MS data from RAW files.[62, 63] Manual validation of more than 20 exemplary spectra revealed that, after conversion of proprietary RAW files to mgf or mzML files by Proteome Discoverer 2.0, ca. 25% of monoisotopic precursor peaks were wrongly assigned. Thus, we implemented an algorithm in Merox 2.0 that analyzes the mass spectrum associated to each MS/MS spectrum in an mzML file to eventually correct its precursor mass (vide supra).

### Supplementary Figures

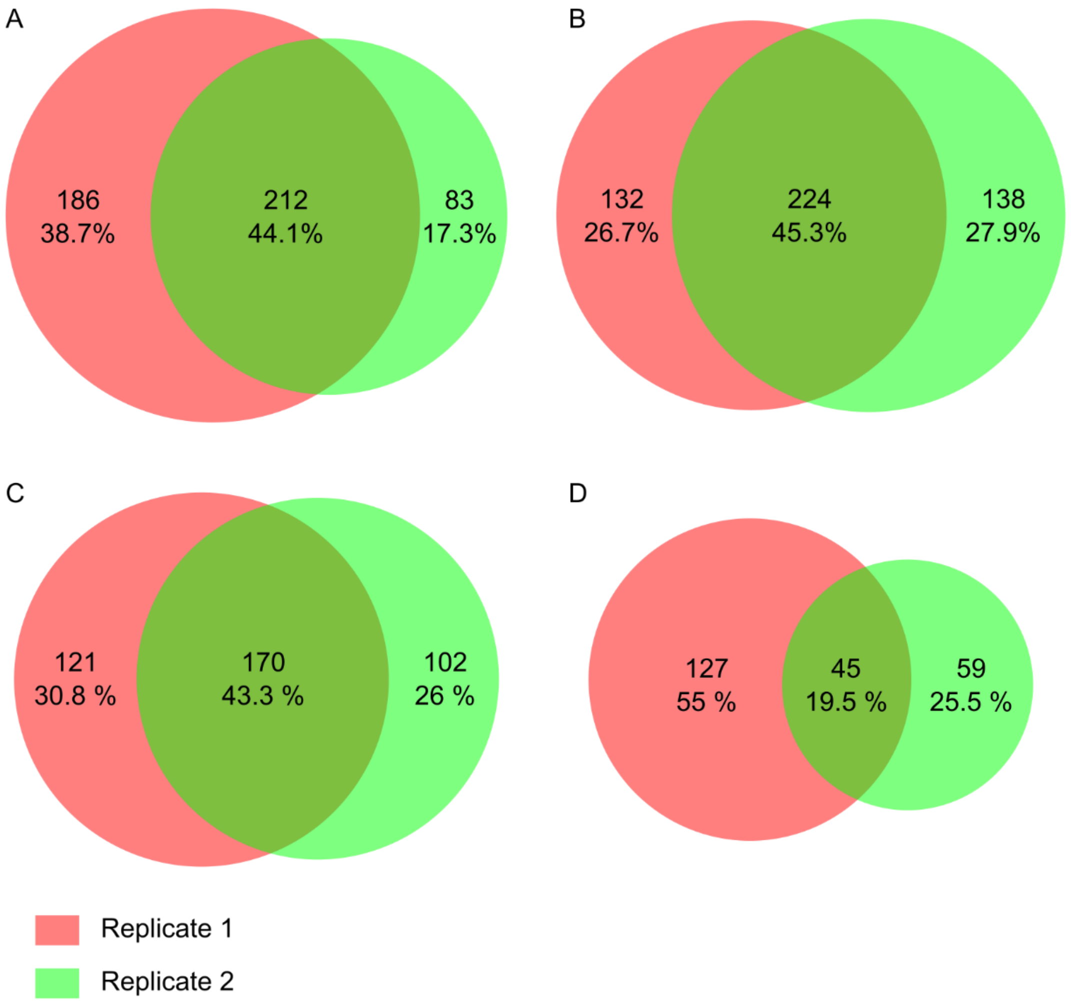


**Figure S1:** Comparison of technical replicates from four SEC fractions. Due to the (i) high complexity of the fractions, (ii) low abundance of cross-linked peptides, and (iii) low excess of cross-linker, the overlap is ~ 45%.

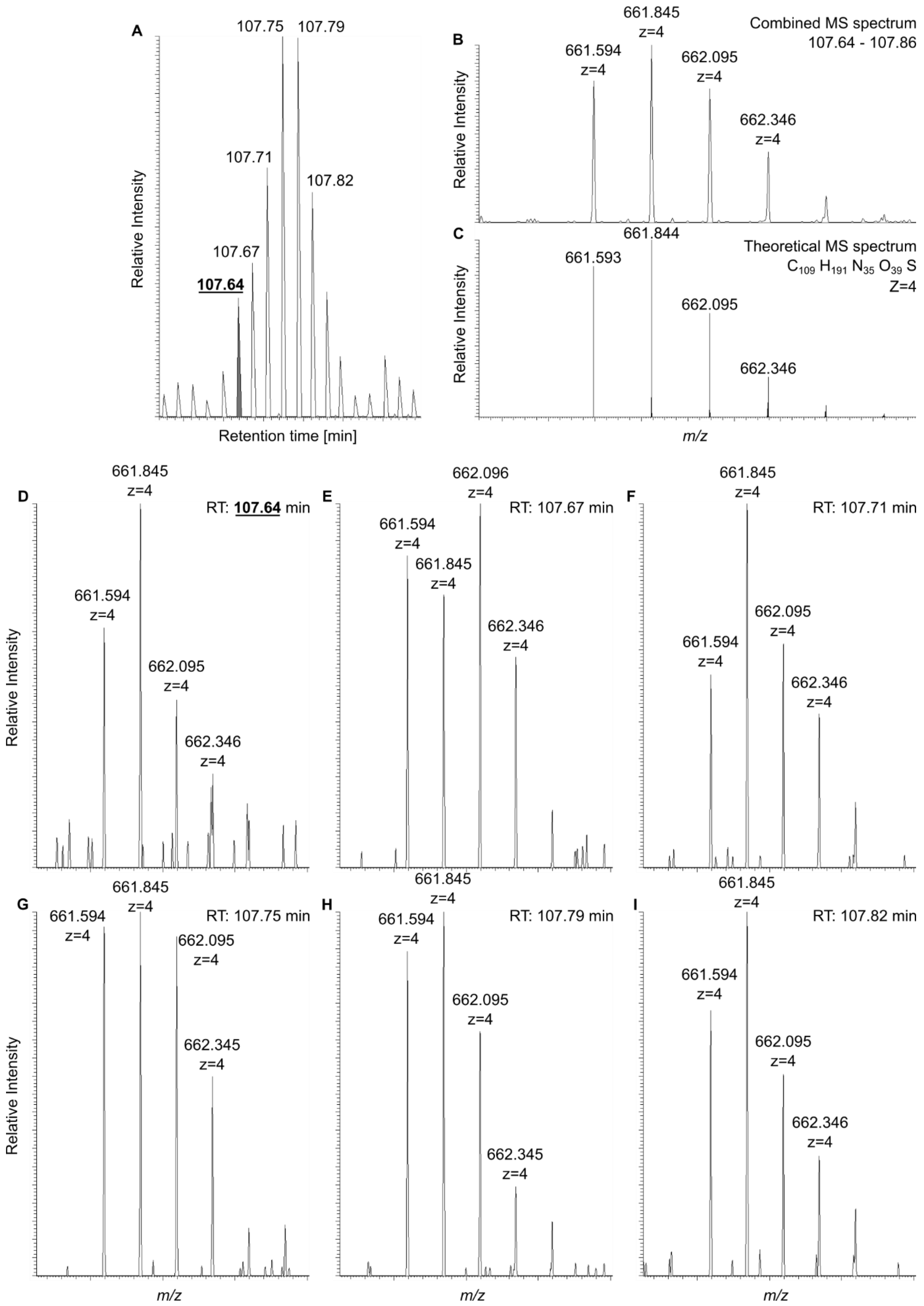
A
107.75 107.79
107.71
107.82
107.67
107.64
Relative Intensity
Retention time [min]
B
Combined MS spectrum
107.64 - 107.86
661.594 z=4
661.845 z=4
662.095 z=4
662.346 z=4
C
Theoretical MS spectrum
$C_{109} H_{191} N_{35} O_{39} S$
Z=4
661.593
661.844
662.095
662.346
m/z
D
RT: 107.64 min
661.845 z=4
661.594 z=4
662.095 z=4
662.346 z=4
E
RT: 107.67 min
662.096 z=4
661.594 z=4
661.845 z=4
662.346 z=4
F
RT: 107.71 min
661.845 z=4
662.095 z=4
661.594 z=4
662.346 z=4
G
RT: 107.75 min
661.594 z=4
661.845 z=4
662.095 z=4
662.345 z=4
H
RT: 107.79 min
661.845 z=4
661.594 z=4
662.095 z=4
662.345 z=4
I
RT: 107.82 min
661.845 z=4
661.594 z=4
662.095 z=4
662.346 z=4
Relative Intensity
m/z

**Figure S2** – Example of monoisotopic peak peaking correction by MeroX 2.0. (A) Extracted ion current of the 4+ charged ion at *m/z* 661.594. The MS/MS event has been triggered between the first two full scans of the corresponding chromatographic peak, at retention times of 107.64 and 107.67 min. (B) Combined full scan mass spectrum in the region of the ion at *m/z* 661.594. (C) Theoretical mass spectrum of the relevant cross-link identified by MeroX 2.0. (D) Full scan mass spectrum at a retention time of 107.46 min. The Excalibur software centered the isolation window on the ion at *m/z* 662.094. During RAW data conversion, the Proteome Discoverer 2.0 assigned the monoisotopic precursor ion to the ion at *m/z* 661.845. MeroX 2.0 applied a "-1" mass correction peaking the ion at *m/*z 661.594. (E-I) Full-scan mass spectra at a retention time range between 107.67 to 107.82 min.

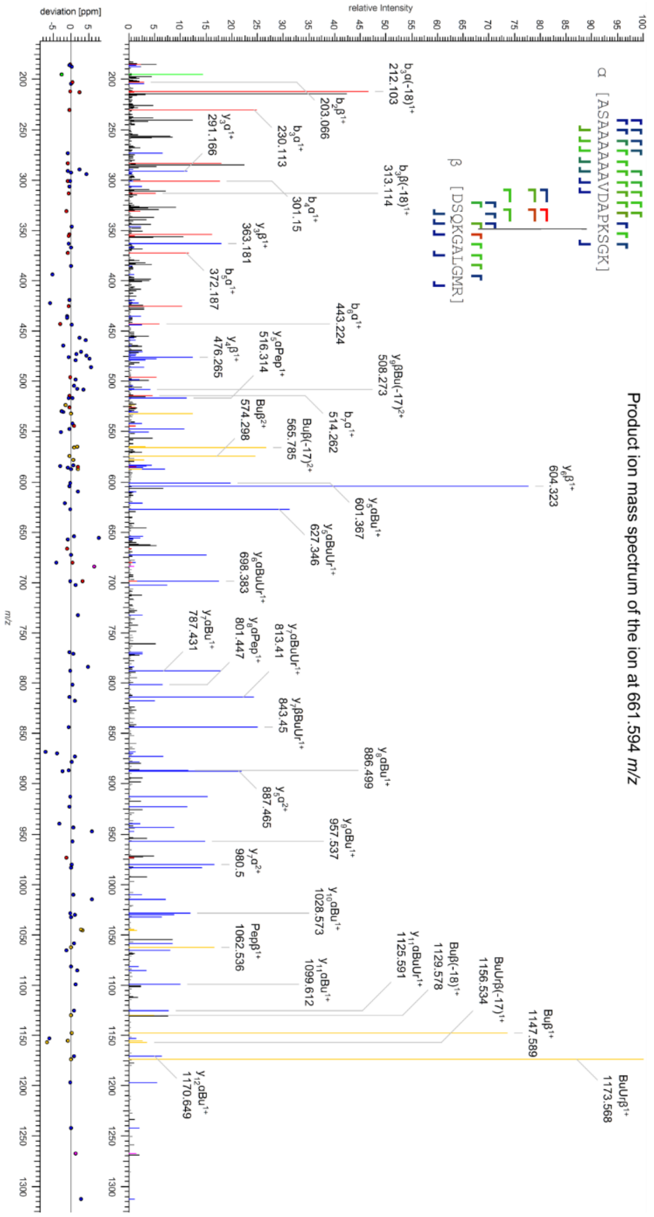


**Figure S3** - Product ion mass spectrum of the 4+ charged ion at *m/z* 661.594. The correct monoisotopic precursor ion was identified by MeroX 2.0 (**Fig. 2**). It was assigned to an intra-protein cross-link between K173 and K182 of eIF3g1. The precursor ion mass deviation is 1.34 ppm.

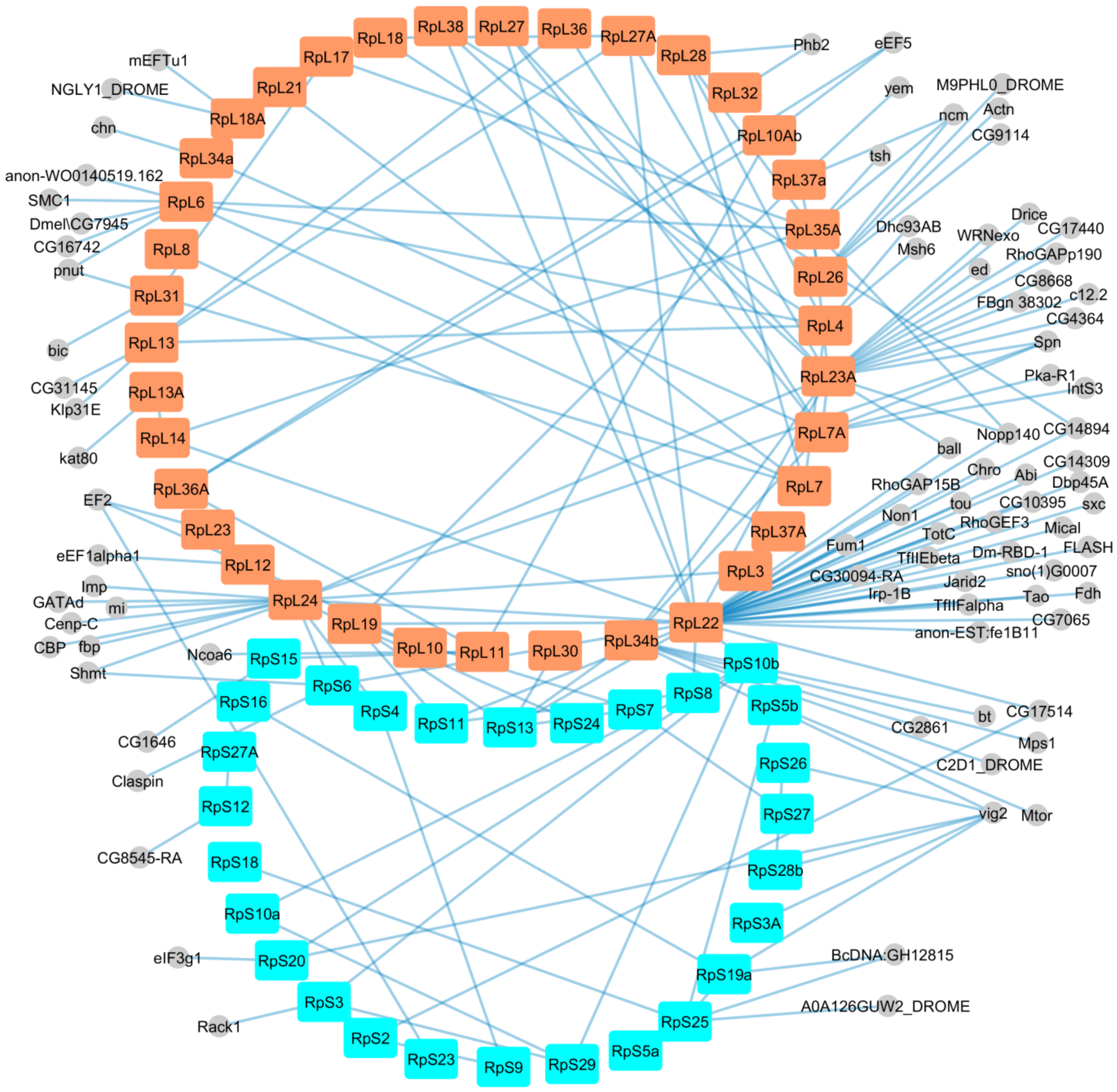


**Figure S4:** Interaction map of 80S ribosome associated proteins. Large subunit proteins are highlighted in orange, small subunit proteins are highlighted in cyan. The figure was prepared using cytoscape.

**A**

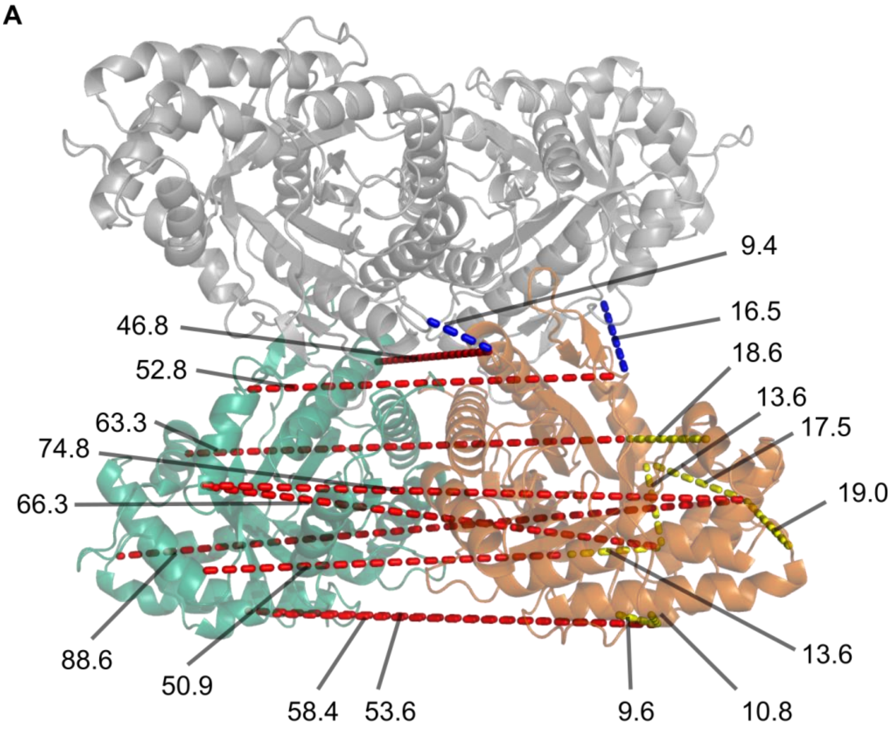


**B**

| | | Cross-links from chain B to: | | | |
|---|---|---|---|---|---|
| **Site 1** | **Site 2** | **Chain B** | **Chain C** | **Chain A** | **Chain D** |
| 42 | 108 | 18.6 | 63.6 | 39 | |
| 156 | 156 | | 46.8 | 46.3 | 9.4 |
| 22 | 99 | 13.6 | 50.9 | 60.8 | |
| 99 | 140 | 13.6 | 66.3 | 48.6 | |
| 111 | 111 | | 52.8 | 16.5 | 52.5 |
| 87 | 317 | 19 | 88.6 | 57.5 | |
| 69 | 329 | 10.8 | 58.4 | 79.1 | |
| 87 | 140 | 17.5 | 74.8 | 42.4 | |
| 69 | 332 | 9.6 | 53.6 | 78.5 | |

**Figure S5:** (A) Cross-links of aldolase were mapped into the X-ray structure of the tetramer of Drosophila aldolase (PDB entry: 2pyo). Two unambiguous inter-protein cross-links are shown in blue. Seven ambiguous cross-links are shown in yellow (intra-protein) and in red (inter-protein). Only ambiguous cross-links between chain C and B are shown in A. All ambiguous cross-links are intra-protein cross-links. Chains B and C of the aldolase homo-tetramer are colored in orange and green, chains A and D are shown in grey. (B) Table of distances (in Å) from chain B to residues in the same or other chains. The color schemes of (A) and (B) are identical.

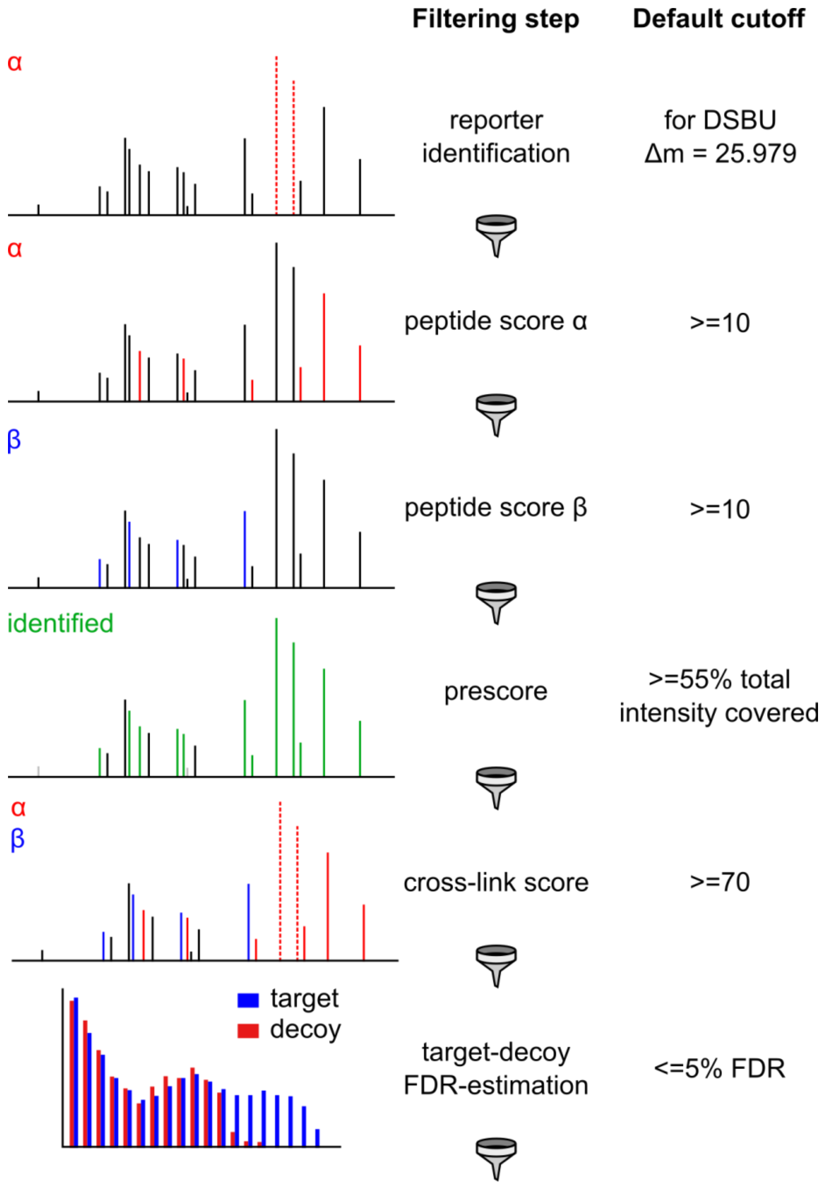


**Figure S6** – Filtering steps during cross-link identification. Spectra are first filtered for the existence of signal pairs with the corresponding cross-link fragment mass difference (25.979 u for DSBU). Peptides are scored against the spectrum to obtain individual peptide scores. Peptides above the specified score cut-off (default 10) will be combined to cross-link candidates that pass through a pre-scoring algorithm where a minimum fraction of signal intensity must be covered by identified ions (default: 55%) to proceed to the scoring algorithm. Only cross-links above the specified score cut-off will be reported (default: 70). After the analysis, candidate hits are filtered by an FDR cut-off that is calculated by parallel analysis of a decoy database.

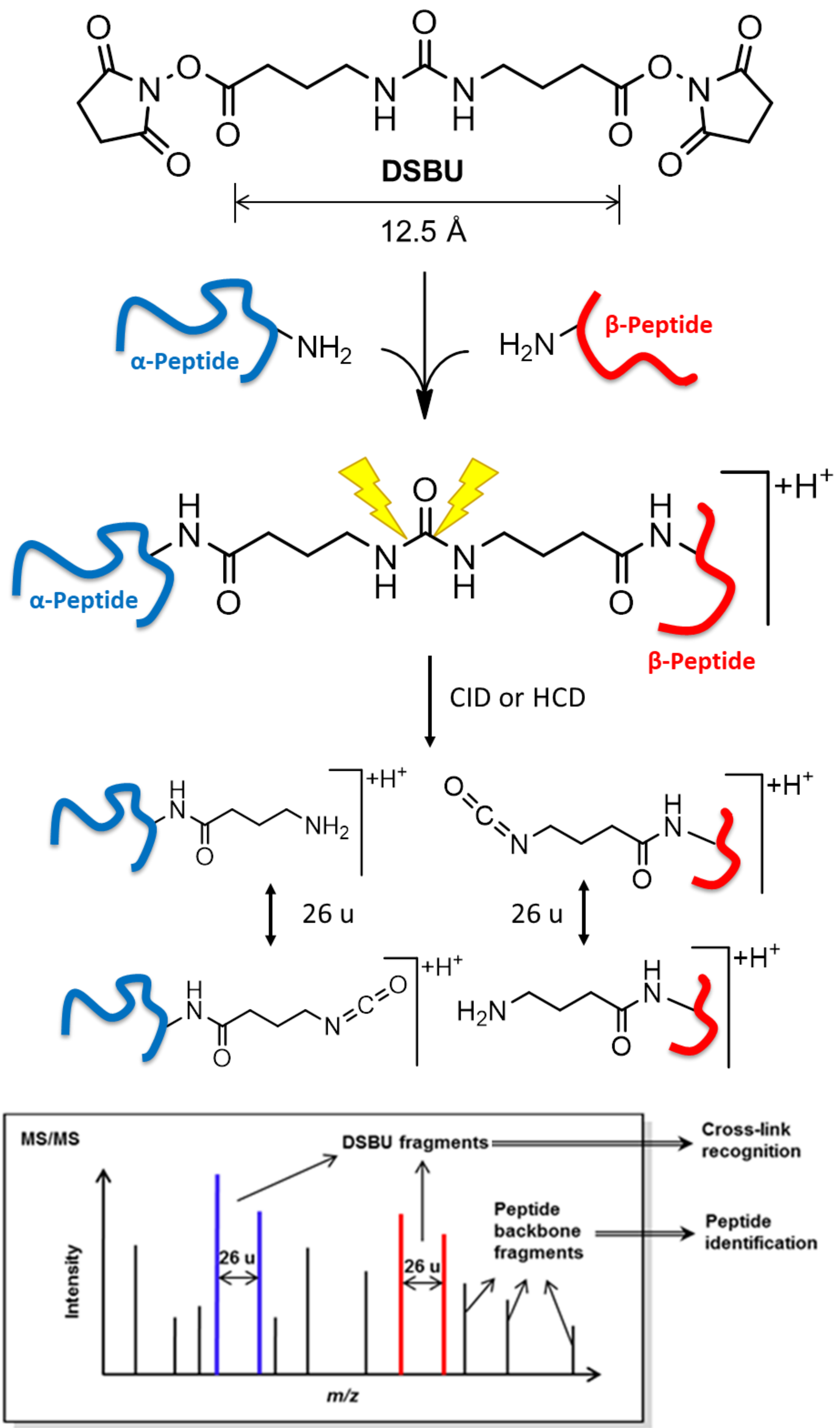
DSBU
12.5 Å
α-Peptide
NH2
H2N
β-Peptide
+H+
CID or HCD
26 u
26 u
MS/MS
DSBU fragments
Cross-link recognition
Peptide backbone fragments
Peptide identification
Intensity
m/z

**Figure S7**: MS-cleavable cross-linker DSBU. DSBU reacts with nucleophilic groups in protein side chains (e.g. primary amines in lysine residues). After proteolytic digest of the cross-linked proteins, two peptides are cross-linked via DSBU. During CID or HCD, the amide bonds of DSBU's urea group are cleaved producing two distinct modifications (+85 u and +111 u) at the cross-linked residue of each peptide ($\Delta m \sim 26$ u). These reporter ions serve to identify cross-linked peptide species and cross-linked peptide masses can be inferred. In addition to the fragmentation at the central urea group of DSBU, peptide backbone fragmentation occurs to give b- and y-type ions. These fragment ions facilitate peptide sequence identification.
